# Parent-of-origin effect *rough endosperm* mutants in maize

**DOI:** 10.1101/054338

**Authors:** Fang Bai, Mary Daliberti, Alyssa Bagadion, Miaoyun Xu, Yubing Li, John Baier, Chi Wah Tseung, Matthew M. S. Evans, A. Mark Settles

**Affiliations:** Horticultural Sciences Department and Plant Molecular and Cellular Biology Program, University of Florida, Gainesville, FL 32611; Biotechnology Research Institute, National Key Facility for Gene Resources and Genetic Improvement, Chinese Academy of Agricultural Sciences, Beijing 100081, China; Department of Plant Biology, Carnegie Institution for Science, Stanford, CA

**Keywords:** parent-of-origin effect, gametophyte, imprinting, seed, endosperm, maize

## Abstract

Parent-of-origin effect loci have non-Mendelian inheritance in which phenotypes are determined by either the maternal or paternal allele alone. In angiosperms, parent-of-origin effects can be caused by loci required for gametophyte development or by imprinted genes needed for seed development. Few parent-of-origin effect loci have been identified in maize (*Zea mays*) even though there are a large number of imprinted genes known from transcriptomics. We screened *rough endosperm* (*rgh*) mutants for parent-of-origin effects using reciprocal crosses with inbred parents. Six *maternal rough endosperm* (*mre*) and three *paternal rough endosperm* (*pre*) mutants were identified with three *mre* loci mapped. When inherited from the female parent, *mre*/+ seeds reduce grain-fill with a rough, etched, or pitted endosperm surface. Pollen transmission of *pre* mutants results in *rgh* endosperm as well as embryo lethality. Eight of the loci had significant distortion from the expected one-to-one ratio for parent-of-origin effects. Linked markers for *mre1*, *mre2*, and *mre3* indicated that the mutant alleles have no bias in transmission. Histological analysis of *mre1*, *mre2*, *mre3*, and *pre*-949* showed altered timing of starch grain accumulation and basal endosperm transfer cell layer (BETL) development. The *mre1* locus delays BETL and starchy endosperm development, while *mre2* and *pre*-949* cause ectopic starchy endosperm differentiation. We conclude that many parent-of-origin effects in maize have incomplete penetrance of kernel phenotypes and that there is a large diversity of endosperm developmental roles for parent-of-origin effect loci.

## INTRODUCTION

The maternal and paternal parents have different genetic and epigenetic contributions to angiosperm seed development. Angiosperm seeds result from the double fertilization of two multicellular gametophytes (WALBOT AND EVANS 2003). In diploid species, gametophytes grow from the haploid products of meiosis with the male and female gametophytes following different developmental programs. The male gametophyte or pollen grain, delivers two haploid sperm cells through the pollen tube to fertilize the female gametophyte. Fertilization of the egg forms a diploid zygote, and fertilization of the two central cell nuclei forms a triploid endosperm cell. The central cell and egg cell provide the vast majority of cytoplasm for the nascent endosperm and the zygote. In addition, the central cell genome has more open chromatin, and there is substantial evidence for a dominant maternal role to initiate the coordinate development of the endosperm and embryo (BAROUX AND AUTRAN 2015; BORG AND BORG 2015; DEL TORO-DE LEON et al. 2016).

Mutations in loci specific to the development of either gametophyte are expected to show non-Mendelian inheritance such as reduced transmission and maternal effect seed phenotypes. Only a few maize seed mutants have been identified with maternal effects, and most of these mutants primarily affect gametophyte development. The *indeterminate gametophyte1* (*ig1*) locus encodes a LATERAL ORGAN BOUNDRIES (LOB) domain transcription factor that is required to limit cell divisions in the female gametophyte (KERMICLE 1971; EVANS 2007). Plants that are heterozygous for *ig1* give a high frequency of defective kernels when pollinated with normal inbred lines. Similarly, *baseless1* (*bsl1*) heterozygous plants will segregate near 1:1 defective kernels when pollinated with inbred pollen (GUTIERREZ-MARCOS et al. 2006). The polar nuclei of the *bsl1* central cell are not positioned correctly in the female gametophyte indicating defective embryo sac development is likely to alter kernel development. The maize *stunter1* (*stt1*) locus shows a low frequency of small kernels when fertilized with normal pollen (PHILLIPS AND EVANS 2011). Mutant *stt1* embryo sacs are reduced in size and appear delayed in development. Both *bsl1* and *stt1* show reduced transmission through the male suggesting additional roles in the development of male gametophytes.

As the seed grows, the endosperm supplies nutrients and signals to promote embryo development (YANG et al. 2008; XING et al. 2013; COSTA et al. 2014). The two maternal copies of the genome in the endosperm create a gene dosage difference with maternal alleles expected to provide twice as much gene product as paternal alleles. Despite these differences in gene dosage, mutations in loci required for seed development typically segregate at ratios consistent with Mendelian recessive mutations (NEUFFER AND SHERIDAN 1980; SCANLON et al. 1994; MCELVER et al. 2001; MCCARTY et al. 2005). This pattern of inheritance indicates that a single dose of a normal allele from pollen is expressed sufficiently for most genes essential for seed development. Detailed analysis of recessive seed mutants in *Arabidopsis* indicates that wild-type paternal alleles are in some cases delayed in expression, as measured by genetic complementation of mutant phenotypes, relative to the maternal allele (DEL TORO-DE LEON et al. 2014). Thus, maternal allele expression can be dominant immediately after fertilization, but most genes required for seed development are supplied by both parents.

By contrast, there are genes that have parent-of-origin specific patterns of seed expression known as imprinting (GEHRING et al. 2011; HSIEH et al. 2011; LUO et al. 2011; WATERS et al. 2011; WOLFF et al. 2011; ZHANG et al. 2011; WATERS et al. 2013; XIN et al. 2013; ZHANG et al. 2014). Imprinted genes are epigenetically regulated such that gene expression is biased as either paternally expressed genes (PEG) or maternally expressed genes (MEG). Like gametophyte mutants, mutations in imprinted loci required for seed development are expected to show non-Mendelian segregation. In *Arabidopsis*, these parent-of-origin effects can manifest as mutants with half seed set, such as 1:1 segregation for defective seeds or aborted ovules. Molecular studies of *Arabidopsis* maternal-effect loci identified the FERTILIZATION INDEPENDENT SEED Polycomb Repressor Complex 2 (FIS-PRC2) as a primary regulator of early endosperm development (OHAD et al. 1996; CHAUDHURY et al. 1997; GROSSNIKLAUS et al. 1998; KIYOSUE et al. 1999; KOHLER et al. 2003). FIS-PRC2 trimethylates lysine 27 on histone H3 to add repressive chromatin marks, which are required for imprinted patterns of gene expression (KOHLER et al. 2012). Mutants in FIS-PRC2 allow central cell divisions prior to fertilization and cause aberrant endosperm and embryo development. Even though most of the *Arabidopsis* FIS-PRC2 subunits have a MEG pattern of gene expression, the primary seed defect results from the loss of the complex in the female gametophyte (LEROY et al. 2007). Mutations in additional *Arabidopsis* MEG and PEG loci have been identified with few showing seed phenotypes (BAI AND SETTLES 2014; WOLFF et al. 2015).

In maize, the *maternally expressed gene1* (*meg1*) is imprinted during the early stages of basal endosperm transfer layer (BETL) development and is expressed from both maternal and paternal alleles later in development (GUTIERREZ-MARCOS et al. 2004). The BETL transfers nutrients from the maternal to filial tissues, and *meg1* encodes a small peptide that promotes differentiation of the BETL (COSTA et al. 2012). Maternal control of *meg1* provides a mechanism to determine the size of the BETL thereby influencing sink strength of individual developing kernels. The *maternal effect lethal1* (*mel1*) locus in maize may also identify a maternal factor that determines grain-fill (EVANS AND KERMICLE 2001). Plants heterozygous for *mel1* show a variable frequency of reduced grain-fill kernels, but these are unlikely to be caused by the female gametophyte as no embryo sac defects are apparent in the mutant. Molecular studies of *mel1* have been limited, because the mutant is only expressed in a single inbred background and requires at least two sporophytic enhancer loci.

Despite being maternal effect loci, both *stt1* and *mel1* can have a frequency of defective kernels well below the 1:1 ratio expected for a parent-of-origin effect locus. Here we report a systematic genetic approach to identify maize parent-of-origin effect loci even with a variable expressivity and low penetrance of seed developmental defects. A screen of 193 defective kernel mutants showing *rough endosperm* (*rgh*) phenotypes identified six maternal-effect and three paternal-effect loci. Mapping of three *mre* mutants indicates that these are new parent-of-origin effect loci with all loci having normal transmission through both male and female gametes. Characterization of the mutant developmental phenotypes reveals that parent-of-origin effect mutants can result in aberrant differentiation of specific endosperm cell types as well as delayed endosperm differentiation.

## MATERIALS AND METHODS

### Genetic stocks

All genetic experiments were completed at the University of Florida Plant Science Research and Education Unit in Citra, FL or greenhouses located at the Horticultural Sciences Department in Gainesville, FL. For the parent-of-origin effect screen, normal seed were planted from segregating self-pollinations of 193 independent *rgh* mutants isolated in the UniformMu transposon-tagging population (MCCARTY et al. 2005). Each mutant isolate was self-pollinated and crossed onto the B73 and Mo17 inbred lines. Pollen from B73 and Mo17 was crossed onto the second ears of mutant isolates when possible. All crosses were screened for *rgh* kernel phenotypes, and the frequency of *rgh* phenotypes were compared between inbred crosses and segregating self-pollinations. Putative *mre* and *pre* mutants were sown in a subsequent generation and reciprocal crosses were completed with the W22 inbred line.

Backcross (BC_1_) mapping populations were developed by crossing F_1_ hybrids with both inbred parental lines. For example, Mo17 x *mre1*/+ F_1_ progeny were crossed reciprocally with Mo17 and W22. BC_1_ ears that segregated for the *mre* phenotype were then used for molecular mapping and transmission analysis.

Mature and developing kernel phenotype analysis was completed with mutant by W22 inbred crosses with plants segregating for *mre* or *pre* genotypes. The *mre/+* X W22 pollinations were dated and sampled; second ears were crossed and scored for *mre* phenotypes at maturity. For W22 X *pre*-949*/+ developmental analysis, plants segregating for *pre*-949*/+ genotypes were crossed onto two W22 plants with one pollination scored for *pre*-949* phenotypes at maturity.

### Mature kernel phenotypes

Segregating *mre/+ and +/pre* crosses with the W22 inbred were visually sorted into mutant and normal sibling kernels. Single-kernel near infrared spectroscopy was used to predict quantitative kernel traits for 96 normal and 96 mutant kernels of each isolate (SPIELBAUER 2009; GUSTIN et al. 2013). Predicted traits include: weight (mg), % oil, % protein, % starch, seed density (g/cm^3^), material density (g/cm^3^), seed volume (mm^3^), and material volume (mm^3^). Sagittal sections of mature kernels were cut with a fixed-blade utility knife and imaged on a flatbed scanner.

### Molecular mapping

BC_1_ progeny from crosses *mre1/+* X Mo17, *mre2/+* X B73, and *mre3/+* X Mo17 were sorted for *rgh* phenotypes. DNA was extracted as described (SETTLES et al. 2004) from individual *rgh* kernels as well as normal sibling pools of 12 kernels per pool. For *mre1*, simple sequence repeat markers (SSRs) were selected from pre-screened SSRs to have one polymorphic marker per chromosome arm (MARTIN et al. 2010). Each marker was amplified from 24 *rgh* kernels and scored for recombination. Segregation distortion was found for umc1294. Two linked markers, umc1164 and phi021, were amplified and scored to determine the region for fine-mapping. For *mre2* and *mre3*, DNA was extracted from 36 BC_1_ *rgh* kernels for each mutant. Each DNA sample was genotyped using the Sequenom MassARRAY platform at the Iowa State University Genomic Technologies Facility as described (LIU et al. 2010b) except that a subset of 144 distributed single nucleotide polymorphism (SNP) markers were genotyped for each sample. Recombination frequencies for each marker were used to identify regions that had significant distortion for fine-mapping. Additional SSR markers and insertion-deletion polymorphism (InDel) markers were screened for the fine-mapping regions on chromosome 4, 6, and 10 as described (SETTLES et al. 2014). DNA was extracted from expanded BC_1_ populations, amplified, and scored for recombination. Primer sequences for SSR and InDel markers are given in Table S2.

### Transmission assay

F_1_ hybrids of *mre1* with Mo17, *mre2* with B73, and *mre3* with Mo17 were reciprocally crossed to generate BC_1_ progeny with heterozygotes as either the male or female parent. The crosses were screened for *mre* phenotypes to select heterozygous F_1_ individuals for transmission analysis. For each cross, 100 BC_1_ kernels were systematically sampled from kernel rows along the tip to base axis of the ear. Transmission of the mutant locus was scored using linked markers proximal and distal to each mutant locus. Primer sequences for the molecular markers are in Table S2.

### Histochemical staining of developing seeds

Developing ears were harvested from 6 DAP to 19 DAP of *mre1/+* X W22, *mre2/+* X W22, *mre3/+* X W22, and W22 X *+/pre*-949*. Harvest dates were adjusted in the spring and fall season due to temperature differences during the June and November kernel development periods. Kernels were fixed in FAA solution (3.7% formaldehyde, 5% glacial acetic acid, and 50% ethanol) at 4°C overnight. Kernels were dehydrated in an ethanol series, and then embedded in paraffin or JB-4 Plastic embedding media (Electron Microscopy Sciences). Paraffin-embedded sample were cut into 8 μm longitudinal sections close to the sagittal plane, deparaffinized, rehydrated, and counterstained with 1% safanin O and 0.5% fast green as described (BAI et al. 2012). Resin embedded samples were cut into 4 μm sections. The sections were treated with 1% periodic acid for 10 minutes, rinsed in the running water for 5 minutes, and then placed in Schiff’s reagent for 30 minutes. The sections were transferred through three successive baths of 2 minutes each of 0.5% sodium metabisulfite in 1% HCl. Sections were then rinsed in running water for 5 minutes, countered stained in 1% aniline blue-black in 7% acetic acid for 20 minutes, rinsed in 7% acetic acid, and rinsed in water. Sections were dried, mounted, and examined by light microscopy. Images were captured with an AmScope digital camera.

### Quantitative RT-PCR

Developing kernels of *mre1/+, mre2/+ and mre3/+* crossed with W22 were sampled at 14 DAP in the fall field season. Kernels were cut in half with a transverse section as described (GOMEZ et al. 2009). Total RNA was extracted from the basal section of the kernel. Briefly, 100 mg of ground tissue was mixed with 200 μL of RNA extraction buffer (50 mM Tris-HCl, pH 8, 150 mM LiCl, 5 mM EDTA, 1% SDS in DEPC treated water). The slurry was then extracted twice with 1:1 phenol:chloroform and once with chloroform at 4°C for 5 minutes for each extraction. The aqueous phase was then extracted with Trizol (Invitrogen) and chloroform. RNA was precipitated from the aqueous fraction using isopropoanol and washed with 70% ethanol. RNA pellets were resuspended in nuclease free water (Sigma) treated with Purelink^Tm^ DNAase (Invitrogen). RNA was then further purified using an RNeasy MinElute Cleanup Kit (Qiagen), and 1 μg total RNA was used to synthesize cDNA with M-MLV reverse transcriptase (Promega). Quantitative RT-PCR used a StepOnePlus real-time PCR machine (Applied Biosystems) with 1X SYBR^®^ Green PCR Master Mix (Applied Biosystems) as described (FOUQUET et al. 2011). The normalized expression level of each gene represents the average of three replicates of three distinct kernel pools relative to *Ubiquitin* using the comparative cycle threshold (*C*_t_) method (LIVAK AND SCHMITTGEN 2001). The primers for each marker gene are listed in Table S2.

### Data and reagent availability

All data necessary for confirming the conclusions are described within the article and supplementary information. Table S2 contains the primer sequences for the molecular markers used in the study. Mutants are available upon request.

## RESULTS AND DISCUSSION

### Parent-of-origin effect screen

We reasoned that parent-of-origin effect mutants with low penetrance could be confused with recessive mutations in large-scale genetic screens, such as the UniformMu genetic screen for defective kernel mutations (MCCARTY et al. 2005). To identify parent-of-origin effects, we reciprocally crossed plants segregating for UniformMu *rough endosperm* (*rgh*) seed phenotypes with B73 and Mo17 inbred pollen. Most *rgh* mutants are seed lethal, and second ears were self-pollinated to identify *rgh* heterozygotes for each isolate. Parent-of-origin effects were distinguished from dominant mutations by comparing self-pollinations to the reciprocal crosses (Figure S1). Mutants were scored as *maternal rough endosperm* (*mre*) if both the self-pollination and cross with inbred pollen segregated for *rgh* phenotypes at similar frequencies, while the *rgh/+* pollen failed to cause seed mutant phenotypes. The *paternal rough endosperm* (*pre*) mutants segregated for *rgh* phenotypes in self-pollinations and crosses onto inbred ears, while crosses of *pre/+* with inbred pollen developed all normal seeds. This strategy requires two ears to be successfully pollinated on individual *rgh/+* plants. A total of 146 *rgh* isolates had sufficient crosses to be screened for both *mre* and *pre* phenotypes. An additional 47 isolates lacked the *rgh/+* by inbred cross and were screened for putative *pre* phenotypes, which could also have been dominant mutations.

Eight putative *mre* and seven putative *pre* isolates were identified and additional reciprocal crosses were completed with the W22 inbred. These crosses showed that six *mre* and three *pre* isolates had consistent parental effects with multiple inbred parents (Figure 1, Figure S2). We found a wide range of segregation ratios for defective kernels in the *mre* and *pre* isolates (Table 1). Only *mre3* had a 1:1 ratio of defective to normal seeds suggesting that *mre* and *pre* loci either have reduced transmission or reduced penetrance of the *rgh* kernel phenotype.

**Figure 1.**
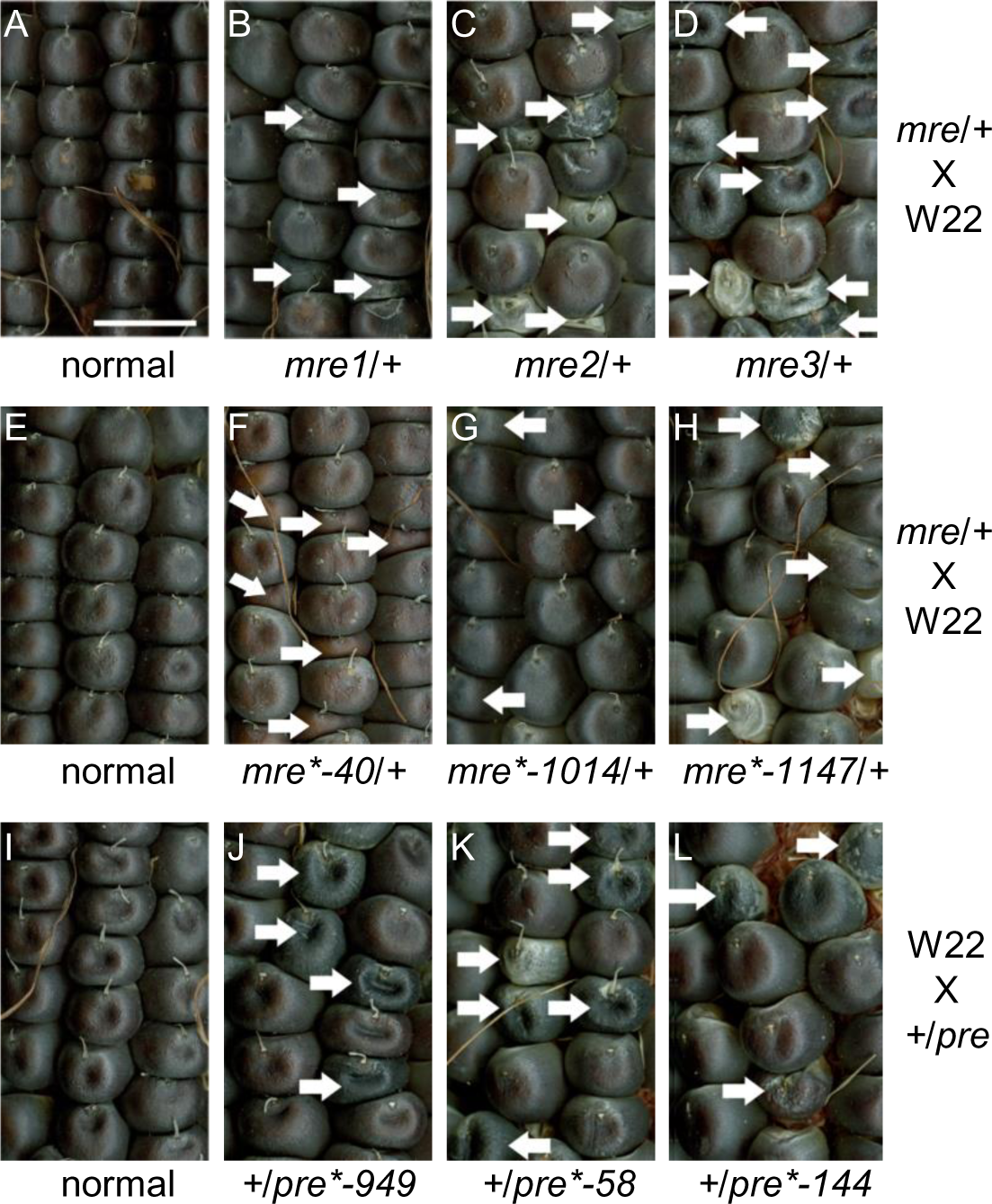
Reciprocal crosses reveal six *mre* mutants and three *pre* mutants. Parent-of-origin effect mutants identified from 193 UniformMu *rgh* isolates. (**A**) normal sibling of *mre1* X W22, (**B**) *mre1/+* X W22, (**C**) *mre2/+* X W22, (**D**) *mre3/+* X W22, (**E**) normal sibling of *mre2* X W22, (**F**) *mre*-40/+* X W22, (**G**) *mre*-1014/+* X W22, (**H**) *mre*-1147*/+ X W22, (**I**) W22 X normal sibling of *pre*-949*, (**J**) W22 X *+/pre*-949,* (**K**) W22 X *+/pre*-58*, and (**L**) W22 X *+/pre*-144*. White arrows indicate mutant seeds. All panels are at the same scale with the bar showing 1 cm in (**A**).

**Table 1.** Segregation of *mre* and *pre* mutants in W22 crosses.

| Cross | Isolate | *rgh* | Normal | % *rgh* | Ratio | $p(\chi^2)$ for 1:1 ratio |
| --- | --- | --- | --- | --- | --- | --- |
| *mre*/+ X W22 | *mre1* | 351 | 436 | 44.6 | 1:1.24 | $2.4 \times 10^{-3}$ |
| | *mre2* | 276 | 450 | 38.0 | 1:1.63 | $1.1 \times 10^{-10}$ |
|  | *mre3* | 302 | 333 | 47.6 | 1:1.10 | 0.22 |
| | *mre* *-40 | 69 | 162 | 29.9 | 1:2.35 | $9.4 \times 10^{-10}$ |
| | *mre* *-1014 | 151 | 280 | 35.0 | 1:1.85 | $5.2 \times 10^{-10}$ |
| | *mre* *-1147 | 25 | 121 | 17.1 | 1:4.84 | $1.9 \times 10^{-15}$ |
| W22 X *pre*/+ | *pre* *-949 | 107 | 348 | 23.5 | 1:3.25 | $1.3 \times 10^{-29}$ |
| | *pre* *-58 | 77 | 194 | 28.4 | 1:2.52 | $1.2 \times 10^{-12}$ |
| | *pre* *-144 | 49 | 136 | 26.5 | 1:2.78 | $1.6 \times 10^{-10}$ |

### Mature seed traits of *mre* and *pre* mutants

Single-kernel near infrared reflectance (NIR) spectroscopy was used to predict kernel composition traits of the *mre* and *pre* isolates (SPIELBAUER 2009; GUSTIN et al. 2013). All mutants reduced seed weight and volume without affecting relative protein and starch content (Figure S4). The three *pre* mutants had significantly reduced oil content, and sagittal sections of mature *pre* mutants revealed embryo development defects (Figure 2A, 2B). Total and material densities were reduced in most of the *mre* and *pre* mutants (Figure S4). Endosperm storage molecule packing influences seed density, and these reductions are consistent with alterations in the mature endosperm such as reduced vitreous endosperm in *mre1* or larger central endosperm air spaces in *mre3* (Figure 2C, 2D).

**Figure 2.**
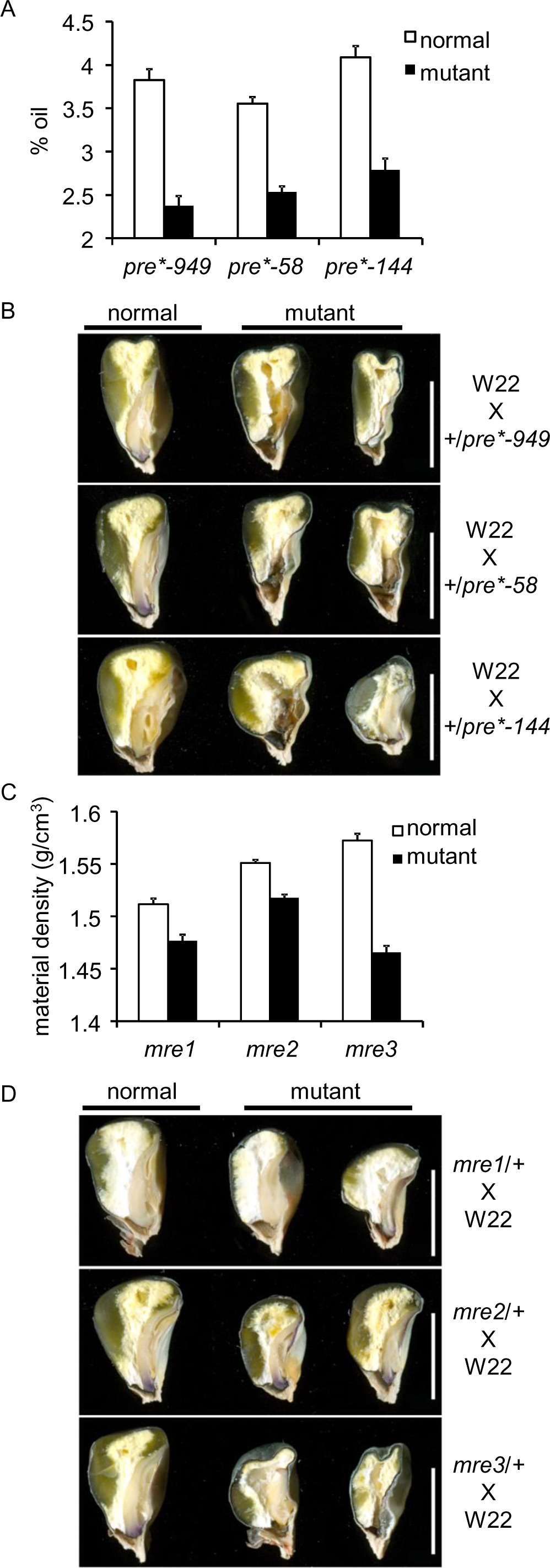
NIR kernel traits and sagittal sections of *mre* and *pre* mature seeds. (**A**) NIR-predicted % oil in *pre*-949*, *pre*-58,* and *pre*-144*. (**B**) Sagittal sections of *pre*-949*, *pre*-58* and *pre*-144*. (**C**) NIR-predicted material density (g/cm^3^) in *mre1*, *mre2,* and *mre3*. (**D**) Sagittal sections of *mre1*, *mre2,* and *mre3*. Scale bar is 0.6 cm in all panels of (**B**) and (**D**).

Sagittal mature kernel sections from *mre* or *pre* mutants showed variable severity in embryo defects suggesting that many of the *mre* or *pre* seeds would fail to germinate (Figure S3, S5). However, oil content was not entirely predictive of *mre* and *pre* mutant germination. Even though *mre1* and *mre2* had no significant reduction in kernel oil content, phenotypically mutant seeds frequently fail to germinate and only a small fraction of the *mre1*/+ and *mre2*/+ seedlings grow and develop normally (Figure 2C, Figure S5). Similarly, *mre*-40* and *mre*-1014* have significantly reduced oil content, yet all mutant seeds germinated with *mre*/+ seedlings being indistinguishable from +/+ siblings (Figure S4, S5). All three *pre* isolates have both low oil and low germination frequency (Figure 2A, Figure S5). These *pre* phenotypes are surprising, because the mutagenic parents for the UniformMu population were crossed as males, and *pre* mutants that fail to germinate would not be expected to survive past the initial mutagenic cross (MCCARTY et al. 2005). All three *pre* isolates have a low frequency of *rgh* kernels when crossed onto inbred ears (Table 1), and it is likely that the *pre* mutants have low penetrance of the mutant phenotype. Both the inheritance patterns and the mature kernel phenotypes of the isolates suggest different developmental mechanisms underlie each *mre* and *pre* mutant phenotype.

### Mapping of *mre1*, *mre2*, and *mre3*

Complementation groups of parent-of-origin effect mutants are not possible to determine with traditional allelism tests. We took a molecular mapping approach to identify specific *mre* and *pre* loci from this screen. F_1_ crosses between each mutant and B73 or Mo17 were then backcrossed to the respective inbred or to the W22 parent of the UniformMu population. These experiments generated BC_1_ backcross mapping populations. For *mre1* and *mre3,* Mo17 was the recurrent mapping parent, and B73 was the recurrent mapping parent for *mre2*. All other isolates failed to segregate for seed phenotypes in any of the BC_1_ crosses. The *mre*-40*, *mre*-1014*, *mre*-1147*, *pre*-58*, *pre*-144*, and *pre*-949* isolates all show *rgh* kernel phenotypes in F_1_ crosses with B73, Mo17, and W22 suggesting allelic differences at each locus, complex genetic modifiers, or epigenetic mechanisms suppress the phenotype of these five mutant loci after crossing to either B73 or Mo17.

To obtain initial map positions, DNA from individual mutant kernels from the BC_1_ populations was genotyped using distributed Simple Sequence Repeat (SSR) or Single Nucleotide Polymorphism (SNP) markers (LIU et al. 2010a; MARTIN et al. 2010). Recombination frequencies were calculated for each marker and the physical position of linked markers identified is listed in Table S1. Expanded mapping populations were scored with additional markers. Figure 3 shows the results of these fine mapping experiments. The *mre1* locus was mapped to a 3.33 Mbp interval on the short arm of chromosome 4, while *mre2* was mapped to a 0.82 Mbp interval on the long arm of chromosome 6. The *mre3* locus maps to a 2.07 Mbp interval on the long arm of chromosome 10 (Figure 3). None of these mutants overlap with the genetic position of published maternal effect mutants including *ig1*, *bsl1*, *stt1*, and *mel1*. These data indicate that *mre1*, *mre2*, and *mre3* are new maternal effect loci.

**Figure 3.**
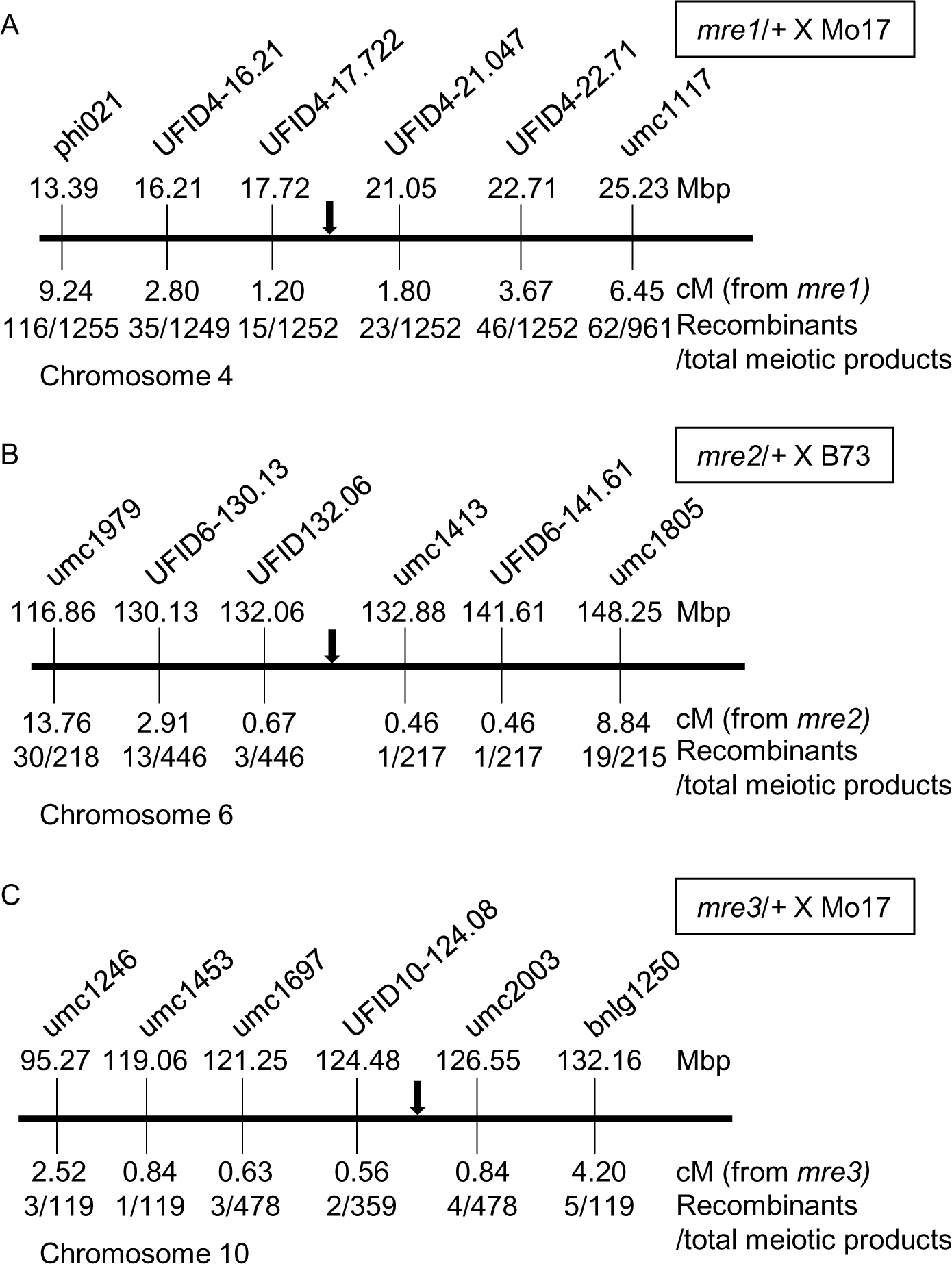
Map positions for three parent-of-origin effect *rgh* loci. Integrated physical-genetic maps for: (**A**) *mre1/+* X Mo17, (**B**) *mre2/+* X B73, and (**C**) *mre3/+* X Mo17 BC_1_ mapping populations. Molecular markers are not positioned to scale. Each schematic indicates chromosome coordinates from the B73_v2 genome assembly for the markers. Recombination frequencies with the mutant phenotypes are given in cM with the number of recombinants and meiotic products scored. The black arrow indicates the mutant locus position.

### Transmission of *mre1*, *mre2*, and *mre3*

The *mre1* and *mre2* loci segregate for less than the 1:1 expected ratio of *rgh* kernels (Table 1), which could indicate incomplete penetrance of the defective kernel phenotype or reduced transmission of the mutant loci. We determined the transmission of each of the mapped loci using linked molecular markers. Reciprocal BC_1_ crosses with heterozygous mutants were sampled along the length of the ear and genotyped with flanking markers (Table S2).

Recombinants between the flanking markers were not included as these kernels could have transmitted either the mutant or normal locus. Ratios close to 1:1 of normal to mutant were observed regardless of the direction of the cross (Table 2). These results indicate that the three *mre* loci transmit fully through both gametes. Based on the frequency of *rgh* kernels in *mre1* and *mre2* crosses, both mutants have incomplete penetrance and a subset of phenotypically normal kernels are expected to be heterozygous for the *mre* loci.

**Table 2.** Transmission of *mre* and *pre* mutant alleles in BC_1_ crosses using linked molecular markers.

| Mutant isolate | Reciprocal cross | W22 (mutant) | Inbred (normal) | Ratio | Expected ratio | Recombinants | $p(\chi^2)$ for 1:1 |
| --- | --- | --- | --- | --- | --- | --- | --- |
| *mre1* | *mre1*/+ X Mo17 | 51 | 44 | 1.16:1 | 1:1 | 5 | 0.47 |
|  | Mo17 X *mre1*/+ | 49 | 46 | 1.07:1 | 1:1 | 5 | 0.76 |
| *mre2* | *mre2*/+ X B73 | 45 | 44 | 1.02:1 | 1:1 | 0 | 0.92 |
|  | B73 X *mre2*/+ | 48 | 50 | 0.96:1 | 1:1 | 2 | 0.84 |
| *mre3* | *mre3*/+ X Mo17 | 53 | 46 | 1.15:1 | 1:1 | 1 | 0.48 |
|  | Mo17 X *mre3*/+ | 56 | 44 | 1.27:1 | 1:1 | 0 | 0.23 |

### Contrasting endosperm defects in *mre3* and *mre1*

It is likely that the *mre* and *pre* mutants disrupt kernel development through different mechanisms. Only the *mre3* mutant is fully penetrant for the mature *rgh* kernel phenotype. We compared endosperm cell morphology in mutant *mre3/+* kernels and normal siblings at two stages of development (Figure 4). The cellularized maize endosperm differentiates into internal starchy endosperm and three epidermal cell fates: aleurone, basal endosperm transfer cell layer (BETL) cells, and embryo surrounding region (ESR) cells (SABELLI AND LARKINS 2009). The starchy endosperm cells in *mre3* mutants are smaller in both developmental stages, but the *mre*/+ cells initiate starch accumulation with similar timing to normal (Figure 4E-F).

**Figure 4.**
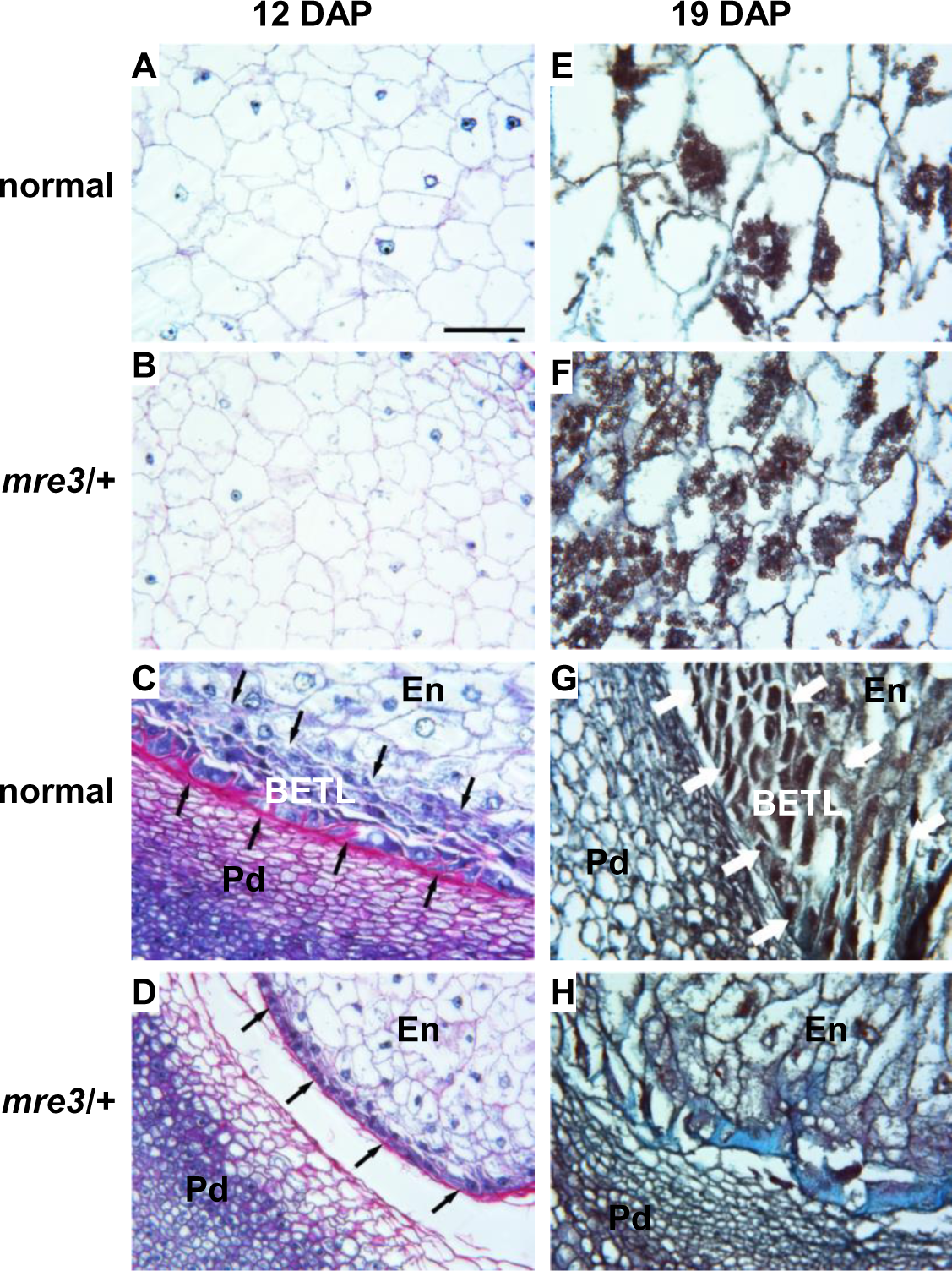
Endosperm defects in *mre3.* (**A-D**) Longitudinal sections of 12 DAP kernels stained with Schiff's reagent and aniline blue-black. Insoluble carbohydrates in cell walls and starch grains stain fuschia; nucleoli, nuclei, and cytoplasm stain different intensities of blue. (**E-H**) Longitudinal sections of 19 DAP kernels stained with safranin and fast green. Starch and secondary cell walls are intensely stained. All samples are collected during the fall season. (**A, E**) Central endosperm of normal sibling kernels. (**B, F**) Central endosperm of *mre3* kernels. (**C, G**) BETL endosperm region of normal kernels. (**D, H**) BETL endosperm region of *mre3* kernels. Arrows indicate BETL. All panels are at the same scale, and the bar in (**A**) is 0.1 mm. En, inner endosperm; Pd, pedicel.

The BETL shows more severe defects in *mre3*/+ kernels. The BETL can be clearly identified in normal sibling kernels as multiple layers of elongated transfer cells with extensive secondary cell wall ingrowths at 12 days after pollination (DAP) and 19 DAP (Figure 4C, 4G). The secondary cell wall ingrowths were not found in the BETL region of *mre3*/+ kernels, and the internal layers of cells in the BETL region expand isotropically to resemble starchy endosperm cells (Figure 4D, 4H). These cellular phenotypes suggest *mre3* causes a specific defect in BETL differentiation and bears some similarity with the maize *bsl1* mutant. BETL cells differentiate in patches of the basal endosperm region in *bsl1* mutants (GUTIERREZ-MARCOS et al. 2006).

Similar comparisons between mutant and normal endosperm show a more global endosperm development defect in *mre1* (Figure 5). The *mre1/+* mutants have a general delay in endosperm development with smaller starchy endosperm cells in all developmental stages. Starchy endosperm cells started to accumulate starch granules at 8 DAP in normal sibling seeds (Figure 5E), but no starch granules formed in mutants by 10 DAP (Figure 5J). Mature *mre1*/+ kernels do eventually accumulate starch, because they have equivalent levels of starch and protein to normal siblings at maturity (Figure S4). The endosperm development delay is more clearly seen in the BETL region. At 6 DAP, normal sibling kernels have two layers of elongated transfer cells with extensive secondary cell wall ingrowths (Figure 5C), while no BETL cells are observed in *mre1/+* mutants (Figure 5D). BETL development is clear in both *mre1*/+ and normal siblings after 8 DAP (Figure 5G-K, 5H-L). These phenotypes are similar to the *stt1* locus, which causes reduced grain-fill through a delay in endosperm growth and differentiation (PHILLIPS AND EVANS 2011).

**Figure 5.**
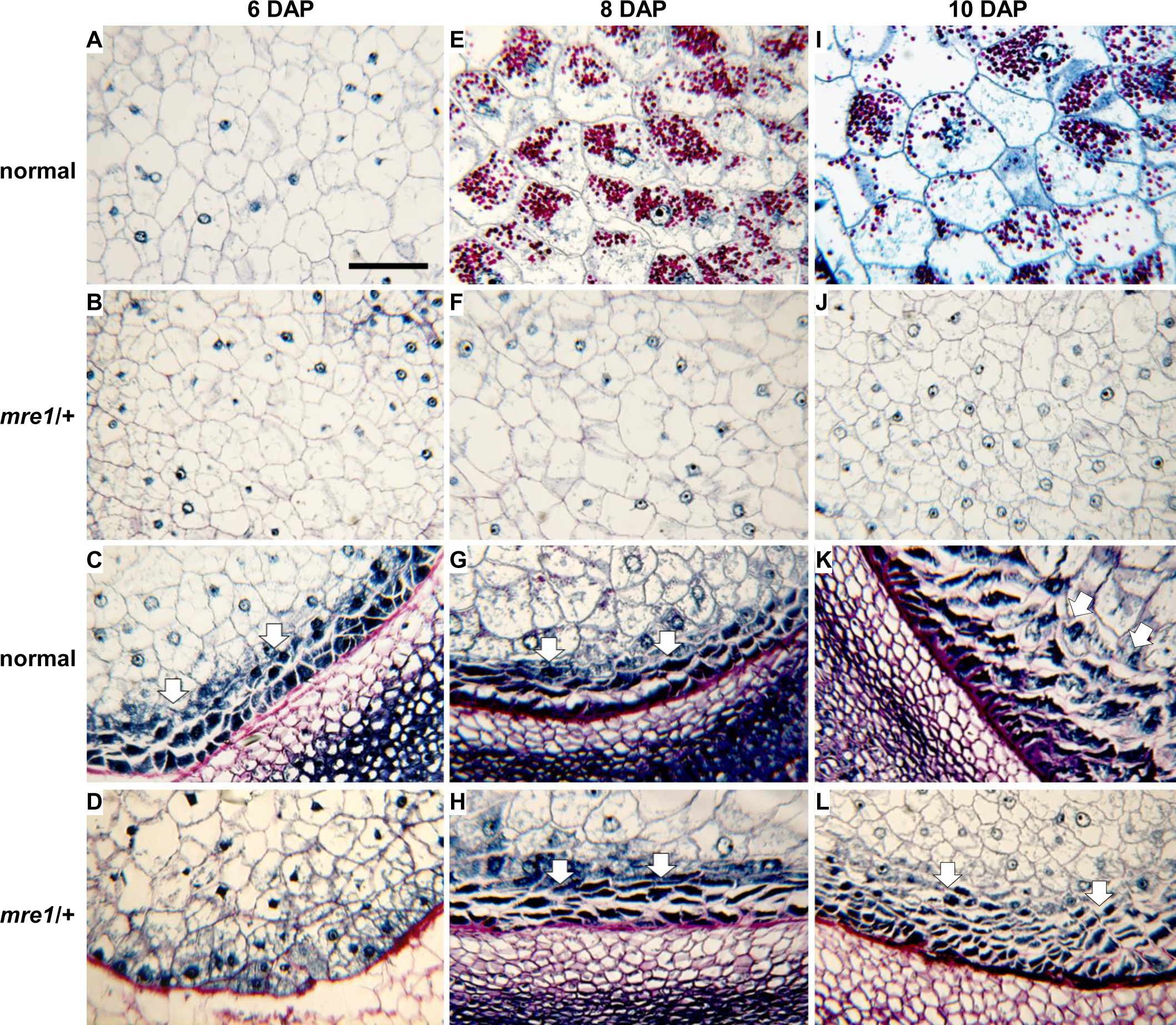
Endosperm development defects in *mre1.* Longitudinal sections through 6 DAP (**A-D**), 8 DAP (**E-H**) and 10 DAP (**I-L**) kernels sampled during the spring field season. All sections were stained with Schiff's reagent and aniline blue-black. Insoluble carbohydrates in cell walls and starch grains stain fuschia; nucleoli, nuclei, and cytoplasm stain different intensities of blue. (**A, E, I**) Central endosperm of normal sibling kernels. (**B, F, J**) Central endosperm of *mre1* kernels. (**C, G, K**) BETL endosperm region of normal kernels. (**D, H, L**) BETL endosperm region of *mre1* kernels. Arrows indicate BETL. All panels are at the same scale, and the bar in (**A**) is 0.1 mm. En, inner endosperm; Pd, pedicel.

We analyzed RNA expression levels of several endosperm cell type markers in *mre1*/+ and *mre3*/+ mutant seeds (Figure 6). Both *Betl2* and *Meg1* are specific to BETL cells, while *Esr1* is specific for ESR cells. The *Rgh3* gene encodes the maize ZRSR2 RNA splicing factor and shows constant expression for the region of the mRNA amplified (FOUQUET et al. 2011). For *mre3*/+, *Betl2* and *Meg1* have large reductions in expression, while *Esr1* is significantly reduced albeit to a lesser extent with about 75% the level of normal kernels (Figure 6A). These data are consistent with a primary *mre3* defect in BETL differentiation. In *mre1*/+ kernels, *Betl2*, *Meg1*, and *Esr1* all have 4-fold or greater reductions, which are consistent with developmental delay of all *mre1* endosperm cell types (Figure 6B).

**Figure 6.**
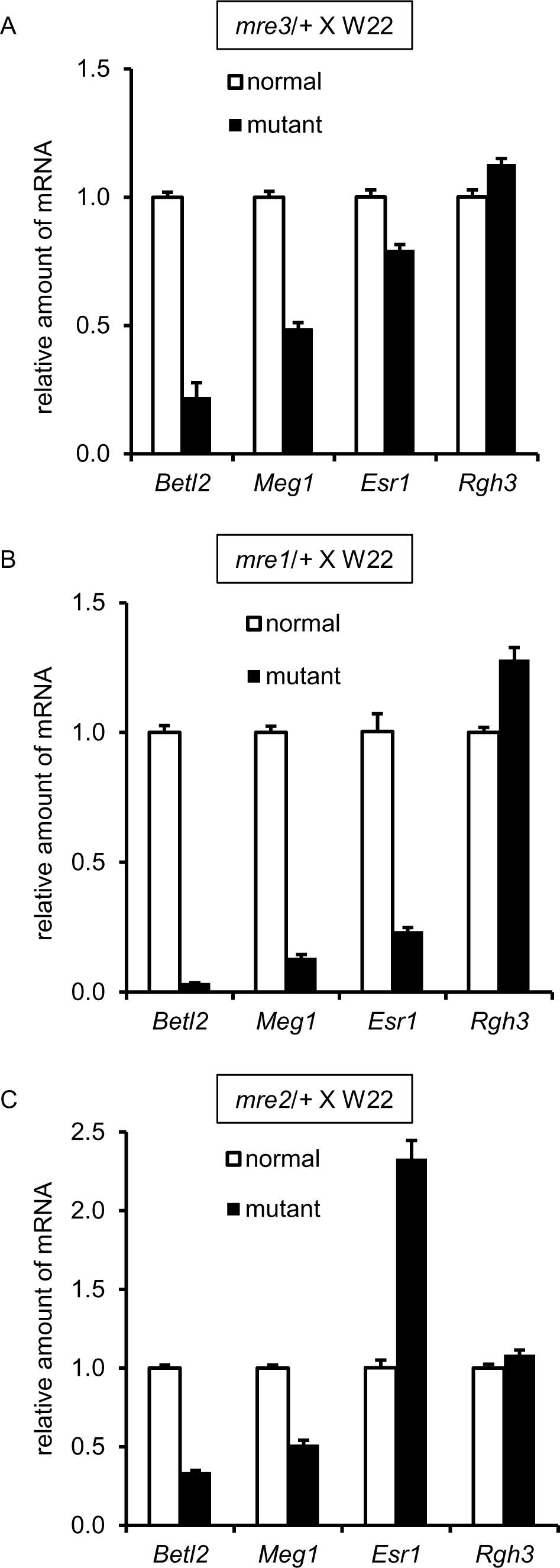
Quantitative RT-PCR of endosperm cell type marker genes in *mre* mutants. Mutant and normal sibling kernels were selected from *mre*/+ X W22 crosses at 14 DAP in the fall season for (**A**) *mre3*, (**B**) *mre1*, (**C**) *mre2*. RNA was extracted from the lower half of the kernels. Values for the *y* axis are arbitrary units of expression level relative to *Ubiquitin*. Error bars indicate standard error of three biological replicates.

### Ectopic endosperm cell differentiation in *mre2* and *pre*-949*

Endosperm cell type marker gene expression in *mre2*/+ kernels showed reductions in *Betl2* and *Meg1*, but more than 2-fold increased *Esr1* expression (Figure 6C). These results indicate that *mre2* confers defects in BETL development and has ectopic *Esr1* expression. Longitudinal sections of developing *mre2*/+ kernels showed multiple cell differentiation defects (Figure 7A-H). In normal seeds, the exterior edge of the endosperm has an epidermal layer and 6-8 starchy endosperm cells with progressive cell expansion towards the center of the endosperm (Figure 7A). The *mre2/+* mutants greatly expanded starchy endosperm cells are found within 2-3 layers of the endosperm epidermal layer (Figure 7E). Starch granules are larger in the *mre2*/+ starchy endosperm cells including in central regions of the endosperm (Figure 7B, 7F). In the BETL region, *mre2*/+ does not develop BETL cells and cells immediately interior to the epidermal layer of the endosperm accumulate starch granules indicating a starchy endosperm cell fate (Figure 7C, 7G). Near the embryo, *mre2*/+ endosperm cells were smaller and without starch granules (Figure 7D, 7H). Combined with *Esr1* expression data, it is likely that *mre2* causes a greater number of ESR cells to differentiate in the endosperm.

**Figure 7.**
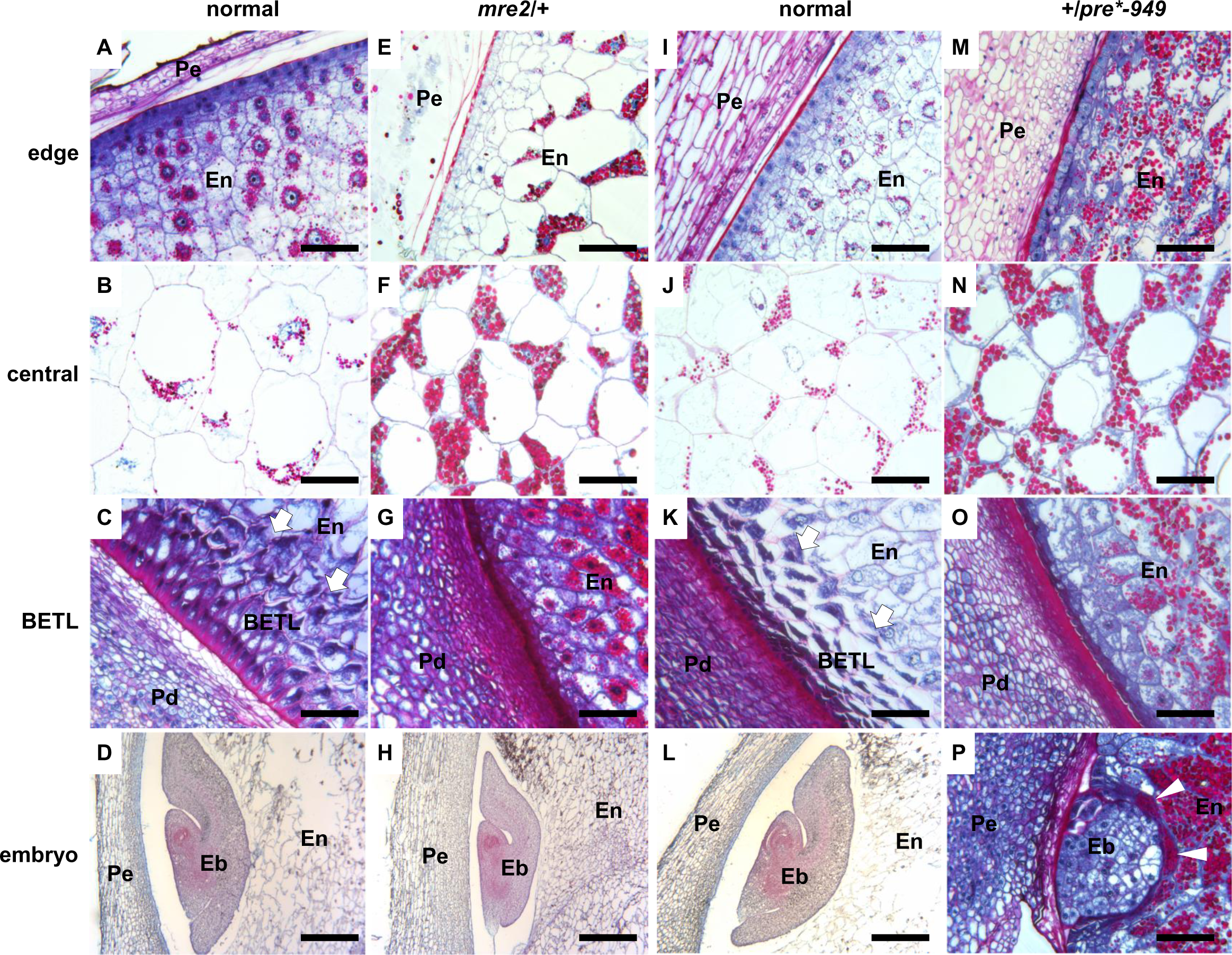
Kernel development defects in *mre2* and *pre*-949*. Longitudinal sections of normal siblings (**AD**, **I-L**), *mre2/+* (**E-H**), and *+/pre*-949* (**M-P**) kernels. Endosperm and the +/*pre*-949* embryo (**P**) sections are stained with Schiff's reagent and aniline blue-black with a 0.1 mm scale bar in each panel. All other embryos (**D, H, L**) are stained with safranin and fast green with a 0.5 mm scale bar in each panel. (**A, E, I, M**) Outer edge of the cellular endosperm (En) and maternal pericarp (Pe). (**B, F, J, N**) Central starchy endosperm. (**C, G, K, O**) Basal endosperm showing the maternal pedicel (Pd), the BETL (arrows), and internal endosperm (En). (D, H, L, P) Maternal pericarp (Pe), embryo (Eb), and endosperm (En). (**A-H**) All sections for *mre2*/+ and normal siblings are from 16 DAP kernels in the fall growing season. (**I-P**) All sections for +/*pre*-949* and normal siblings are from 19 DAP kernels in the fall growing season.

Surprisingly, sections of *+/pre*-949* mutant kernels showed similar endosperm development defects as in *mre2*. The +/*pre*-949* mutants had expanded starchy endosperm cells with starch granules within 1-3 layers of the endosperm epidermis (Figure 7I, 7M). Starch granules are significantly larger in mutants in the central starchy endosperm (Figure 7J, 7N). Moreover, +/*pre*-949* kernels had defective BETL development with the internal cells differentiating into starchy endosperm like in *mre2*/+ mutants (Figure 7K, 7O).

However, +/*pre3*-949* and *mre2/+* show contrasting phenotypes in the ESR region. The +/*pre*-949* ESR differentiates into starchy endosperm and accumulates large starch granules around the embryo, which is arrested at the globular stage (Figure 7L, 7P). Mutant *mre2/+* embryos are smaller but normal in morphology with an enlarged ESR domain (Figure 7D, 7H). The ESR expresses numerous small peptides of the CLE gene family, which are likely involved in cell-to-cell signaling (OPSAHL-FERSTAD et al. 1997; BONELLO et al. 2002; BALANDIN et al. 2005). Moreover, ESR cell differentiation defects are associated with embryo development defects in the maize *rgh3* mutant (FOUQUET et al. 2011). In *Arabidopsis*, the EMBRYO SURROUNDING FACTOR1 (ESF1) gene family is required for normal embryo development and is expressed in the micropylar endosperm (COSTA et al. 2014). Endosperm expression of ESF1 promotes suspensor cell growth and normal basal development in the embryo proper indicating an important role for ESR-like endosperm domains in angiosperm embryo development. Thus, it is likely that ectopic starchy cell differentiation in +/*pre*-949* kernels leads to aborted embryo development.

### Conclusions

Our screen for *mre* and *pre* mutants has revealed that many parent-of-origin effect loci show reduced penetrance of defective kernel phenotypes. These results helps explain the low number of mutant isolates segregating for 50% defective kernels in large-scale genetic screens (NEUFFER AND SHERIDAN 1980; MCCARTY et al. 2005). Phenotyping of reciprocal crosses with inbred lines appears to be a robust method to identify parent-of-origin effect kernel mutants in maize.

The *mre* and *pre* endosperm defects suggest several developmental mechanisms that can give rise to parent-of-origin kernel defects. Defective or delayed BETL cell differentiation was observed in all mutants. The BETL transfers nutrients to the developing seed, and transfer cell defects are likely to limit grain-fill. BETL defects appear to be the primary cause of reduced grain-fill in *mre3* and the *bsl1* loci (GUTIERREZ-MARCOS et al. 2006). A more general delay in endosperm differentiation was found for *mre1*, which is similar to the *stt1* locus (PHILLIPS AND EVANS 2011). By contrast, multiple endosperm cell differentiation defects were found in *mre2* and *pre*-949* with *pre*-949* illustrating the importance of the ESR for maize embryo development. Even though *mre3* and *mre1* have some similarity to *bsl1* and *stt1*, these new loci show no bias in transmission. These data indicate that the female gametophyte is fully functional in the *mre* loci. We believe the most parsimonious explanation for the maternal effects of *mre1*, *mre2*, and *mre3* is that these mutants either encode imprinted, maternally expressed genes. It is also possible that the gene products are stored in the female gametophyte for later seed development functions, or that the *mre* endosperm phenotypes result from interactions between the *mre* female gametophyte and *mre*/+ endosperm. Molecular cloning of the *mre* loci would resolve these alternate models.

## ACKNOWLEDGMENTS

We thank Wei Wu and Mitzi Wilkening at the Iowa State University Genomic Technologies Facility for genotyping services. This work is supported by National Science Foundation (awards IOS-1031416 and MCB-1412218) and the National Institute of Food and Agriculture (awards 2010-04228 and 2011-67013-30032).

## SUPPLEMENTAL MATERIAL

**Figure S1:** Genetic screen for *mre* and *pre* mutants. (**A**) Schematic of pollinations used to screen for parent-of-origin effect mutants. Self-pollination identified plants heterozygous for *rgh* mutations. Reciprocal crosses with inbred lines were screened for *rgh* kernels in the F_1_ generation. (**B**) Self-pollination of *mre1/+* segregates for *rgh* kernels. (**C**) *mre1/+* crossed with Mo17 pollen segregates for *rgh* kernels. (**D**) B73 crossed with *mre1/+* pollen has all normal kernels. Arrows indicate *rgh* kernels.

**Figure S2:** Abgerminal kernel phenotypes of *mre* and *pre* mutants with normal siblings. The six *mre* mutants were crossed with W22 pollen. Pollen from the three *pre* mutants were crossed onto W22 ears. Scale bar is 0.6 cm in all panels.

**Figure S3:** Sagittal sections of mature *mre* and *pre* mutants compared with normal siblings. The six *mre* mutants were crossed with W22 pollen. Pollen from the three *pre* mutants were crossed onto W22 ears. The *mre1*, *mre2*, *mre*-40* and *mre*-1014* mutants frequently develop embryos with shoot and root axes. The *mre3*, *mre*-1147,* and three *pre* mutants frequently are embryo lethal. Scale bar is 0.6 cm in all panels.

**Figure S4:** Single-kernel NIR spectroscopy analysis of kernel traits for the *mre* and *pre* mutants. Spectra were collected from mutant and normal siblings of W22 crosses with heterozygous plants. Mean and standard deviation error bars are plotted for normal siblings (white bars) and mutants (black bars). (**A**) Seed weight (mg/kernel). (**B**) % oil. (**C**) % protein. (**D**) % starch. (**E**) Total seed density (g/cm^3^) including air space. (**F**) Material density (g/cm^3^). (**G**) Total seed volume (mm^3^) including air space. (**H**) Material volume (mm^3^).

**Figure S5:** Germination and seedling phenotypes of a subset of *mre* and *pre* mutants from W22 crosses. Normal and mutant siblings are shown in each panel at 7-8 days after planting. (**A**) *mre1*, (**B**) *mre2,* (**C**) *mre3,* (**D**) *mre*-40,* (**E**) *mre*-1014*, (**F**) *mre*-1147*, (**G**) *pre*-949,* (**H**) *pre*-58,* (**I**) *pre*-144*. Scale bars are 4 cm in all panels.

**Table S1:** Linked markers identified from genome-wide screens.

**Table S2:** Primers for molecular markers used in this study.

